# Regional differences of the urinary proteomes in healthy Chinese individuals

**DOI:** 10.1101/190710

**Authors:** Jianqiang Wu, Weiwei Qin, Li Pan, Fanshuang Zhang, Xiaorong Wang, Biao Zhang, Guangliang Shan, Youhe Gao

**Affiliations:** Department of Pathophysiology, Institute of Basic Medical Sciences, Chinese Academy of Medical Sciences, School of Basic Medicine, Peking Union Medical College, No.5 Dongdansantiao, 100005, Beijing, China; Department of Biochemistry and Molecular Biology, Gene Engineering and Biotechnology Beijing Key Laboratory, Beijing Normal University, No.19 Xinjiekouwai Street, 100875, Beijing, China; Department of Epidemiology and Statistics, Institute of Basic Medical Sciences, Chinese Academy of Medical Sciences, School of Basic Medicine, Peking Union Medical College, No.5 Dongdansantiao, 100005, Beijing, China

**Keywords:** urine, proteomics, Urimem, biobank, biomarkers, regional differences, confounding factors

## Abstract

Urine is a promising biomarker source for clinical proteomics studies. Although regional physiological differences are common in multi-center clinical studies, the presence of significant differences in the urinary proteomes of individuals from different regions remains unknown. In this study, morning urine samples were collected from healthy urban residents in three regions of China and urinary proteins were preserved using a membrane-based method (Urimem). The urine proteomes of 27 normal samples were analyzed using LC-MS/MS and compared among the three regions. We identified 1,898 proteins from Urimem samples using label-free proteome quantification, of which 62 urine proteins were differentially expressed among the three regions. Hierarchical clustering analysis showed that inter-regional differences caused less significant changes in the urine proteome than inter-sex differences. Of the 62 differentially expressed proteins, 10 have been reported to be disease biomarkers in previous clinical studies. Urimem facilitates urinary protein storage for large-scale urine sample collection, and thus accelerates biobank development and urine biomarker studies employing proteomics approaches. Regional differences are a confounding factor influencing the urine proteome and should be considered in future multi-center biomarker studies.

## Introduction

Biomarkers are measurable changes associated with specific pathophysiological conditions. Unlike blood, which is controlled by homeostatic mechanisms, urine reflects systemic changes in the body and can be a more promising source of biomarkers (Gao, 2013; Wu and Gao, 2015). Currently, in addition to its application in urogenital diseases, urinary proteomics has been used in systematic diseases, such as cancers, cardiovascular diseases, diabetes, and mental disorders, to identify urinary disease biomarkers (Shao et al., 2011). More importantly, urine has potential uses in detecting small and early pathological changes in the very early stages of disease (Wu et al., 2017a; Wu et al., 2017b; Yin et al., 2017; Yuan et al., 2015; Zhang et al., 2017). For example, several urinary proteins change significantly even before a tumor mass is palpable, indicating that urine is an excellent source of biomarkers for early cancer detection (Wu et al., 2017a). Another study suggested that early changes in urinary proteins can be detected in a lung fibrosis model, which may lead to early treatment and a better prognosis (Wu et al., 2017b).

In the last decade, advances in proteomics techniques have accelerated the pace of biomarker research. Urinary proteome changes, some of which have been reported to be associated with disease, have been found in various systematic diseases both in animal models and patients. However, candidate biomarkers or biomarker panels need to be validated in large-scale, multi-center patient cohorts. Previous studies showed that several factors can influence the urine proteome (Bakun et al., 2014; Castagna et al., 2011; Kohler et al., 2010; Wu and Gao, 2015; Zhao et al., 2016). Regional differences are a common phenomenon in multi-center clinical studies. These differences may consist of genetic background, food consumption, lifestyle and environmental factors. However, whether there are significant differences in the urinary proteome among different regions remains unknown.

In this study, urine samples were collected from healthy Han residents in three urban areas of China (Haikou, Xi’an and Xining). These three cities are the capitals of Hainan, Shaanxi and Qinghai Provinces, respectively. Urimem is a product that can store urine proteins simply and economically, enabling the large-scale storage of clinical samples (Jia et al., 2014). In the current study, a large number of urine samples were collected at sites in the three regions, and urine proteins were stored using Urimem. Twenty-seven normal Urimem samples were selected and compared based on region to investigate the regional differences in the urinary proteome. Using label-free proteome quantification, a total of 1,898 urine proteins were identified from Urimem samples. A total of 62 urine proteins were differentially expressed among the three regions, of which 10 have been reported to be disease biomarkers. Our findings suggested that regional differences should be taken into consideration in biomarker identification and validation, particularly in multi-region and multi-center clinical studies. This study will not only benefit biobank development but also improve our understanding of biomarker discovery.

## Results

### Characteristics of the participants from the three regions of China

In this study, we recruited healthy urban residents from the capitals of three Chinese provinces (Hainan, Shaanxi, and Qinghai). The geographic distribution of these three regions is shown in **Figure 1**. Haikou is the capital of Hainan Province, located in southern China (longitude: 110.35, latitude: 20.02, elevation: 8 m); Xi’an is the capital of Shaanxi Province, located in central China (longitude: 108.95, latitude: 34.27, elevation: 415 m); and Xining is the capital of Qinghai Province, located in northwest China (longitude: 101.74, latitude: 36.56, elevation: 2250 m). The latitudes of Xi’an and Xining are similar but significantly higher than that of Haikou. The elevations of these three cities are significantly different.

**Figure 1.**
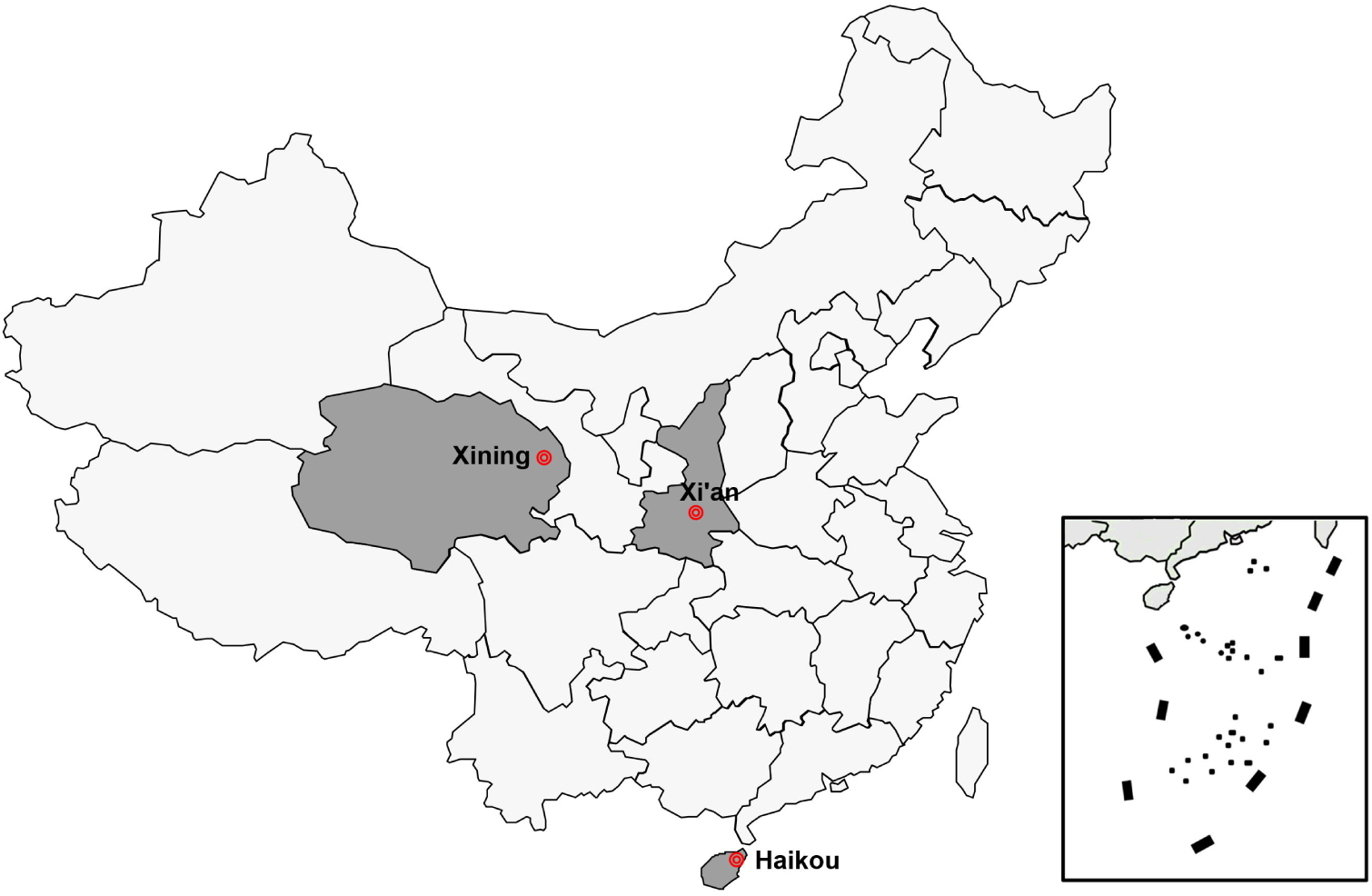
Geographical distribution of the participants enrolled in this study. Xining (longitude: 101.74, latitude: 36.56, elevation: 2250 m); Xi’an (longitude: 108.95, latitude: 34.27, elevation: 415 m); Haikou (longitude: 110.35, latitude: 20.02, elevation: 8 m).

To avoid potential interference from confounding factors, 27 healthy urban Han Chinese adults between 30 and 36 years of age with no chronic or metabolic diseases and no drug treatment within 2 the past weeks were randomly selected. Fasting blood samples were used for laboratory biochemical analyses. Detailed information on the 27 participants is listed in **Supplemental Table S1.** Age, sex, blood pressure and most blood biochemical indicators were similar among the healthy participants from the three areas. However, total protein (TP), albumin (ALB), total cholesterol (TC), low-density lipoprotein cholesterol (LDL-C), fasting venous blood glucose (glucose level) and calcium (Ca) were significantly different (p < 0.05) in the blood of the participants from these three cities (**Figure 2**). Serum protein concentrations, including TP and ALB, were lower in Xi’an residents than in residents from the other cities. The concentrations of serum lipids, including TC and LDL-C, were higher in the Haikou residents than in residents from the other cities. Additionally, glucose level in Xining residents was higher than in residents from Haikou. Meanwhile, serum concentration of Ca was significantly lower in residents from Xi’an than in residents from the other cities.

**Figure 2.**
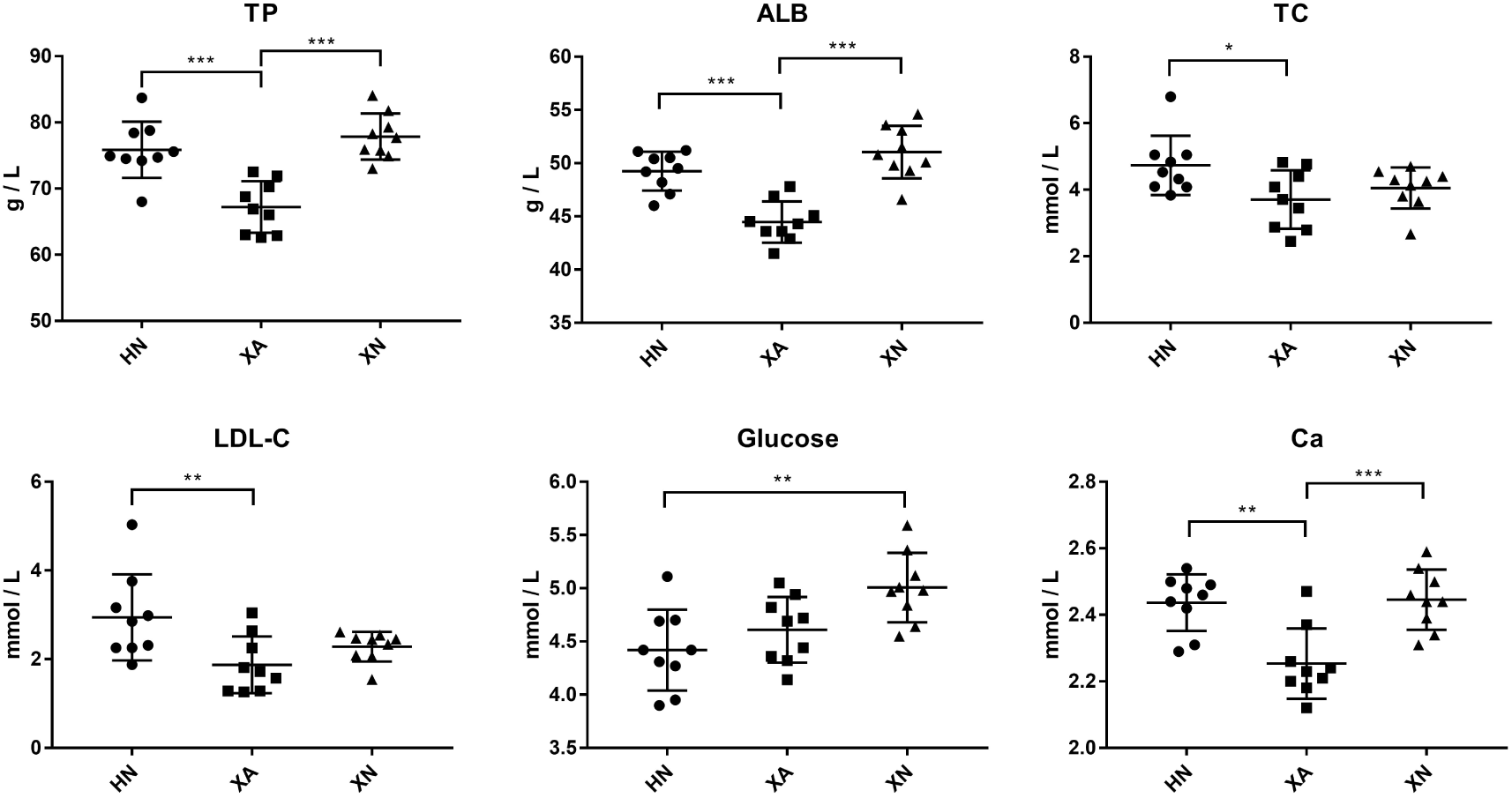
Blood biochemical indicators that differ among three areas of China. Three areas in China: HN (Haikou City), XA (Xi’an City), and XN (Xining City). Abbreviations: TC (serum total cholesterol), LDL-C (low-density lipoprotein cholesterol), Urea (serum urea nitrogen), IgG (Immunoglobulin G), TP (serum total protein), ALB (serum albumin), Ca (calcium) and P (phosphorus). Comparisons between independent regions were conducted using a one-way ANOVA followed by post hoc analysis with the least significant differences test. *indicates p < 0.05, **indicates p < 0.01, and ***indicates p < 0.001.

### Urimem for proteomics analysis

The system of enriching and preserving urinary proteins on membranes using a vacuum filter bottle was first described in our previous study (Jia et al., 2014). Nitrocellulose (NC) membranes were used to absorb the urine proteins in the current study because their reproducibility is similar to protein isolation, and proteins can be easily eluted from them (Wang et al., 2014b). A total of 27 Urimem samples (9 samples from each geographic area) were randomly selected from healthy participants for proteome analysis. During the protein elution, acetone completely dissolved the NC membranes and precipitated the proteins at a low temperature. Urinary proteins were eluted from Urimem, digested with trypsin and then analyzed using one-hour LC-MS/MS. For quantification, Progenesis LC-MS software was used to process the MS data (ion detection, accurate mass retention time pair clustering, normalization and statistically analysis). Finally, a total of 1,898 urine proteins were identified from the Urimem samples using label-free identification (**Supplemental Table S2**).

### Regional differences in the normal human urine proteome

To investigate the regional differences in the normal urine proteome, the Urimem samples from 27 healthy residents from three Chinese regions were analyzed by LC-MS/MS. The participants were matched by age, sex and ethnicity. A total of 62 differentially expressed proteins were identified among the three regions (≥ 2 unique peptides, ANOVA p < 0.05, and fold change > 2). Proteins significantly changed among the three areas were further subjected to the following t-test analyses: Xi’an vs. Haikou, Xi’an vs. Xining and Haikou vs. Xining. There were 43 differential proteins between Xi’an and Haikou, 33 between Xi’an and Xining, and 6 between Haikou and Xining (**Supplemental Table S3**). Hierarchical clustering analysis of the differential proteins (ANOVA p < 0.05) also showed that samples from Xi’an were different from samples in the other two areas, and samples from Haikou and Xining could not be distinguished from one another (**Figure S1**). As shown in **Figure 3A**, there were 59 urine proteins that were differentially expressed between Xi’an and the other cities, 47 between Haikou and the other cities, and 38 between Xining and the other cities.

**Figure 3.**
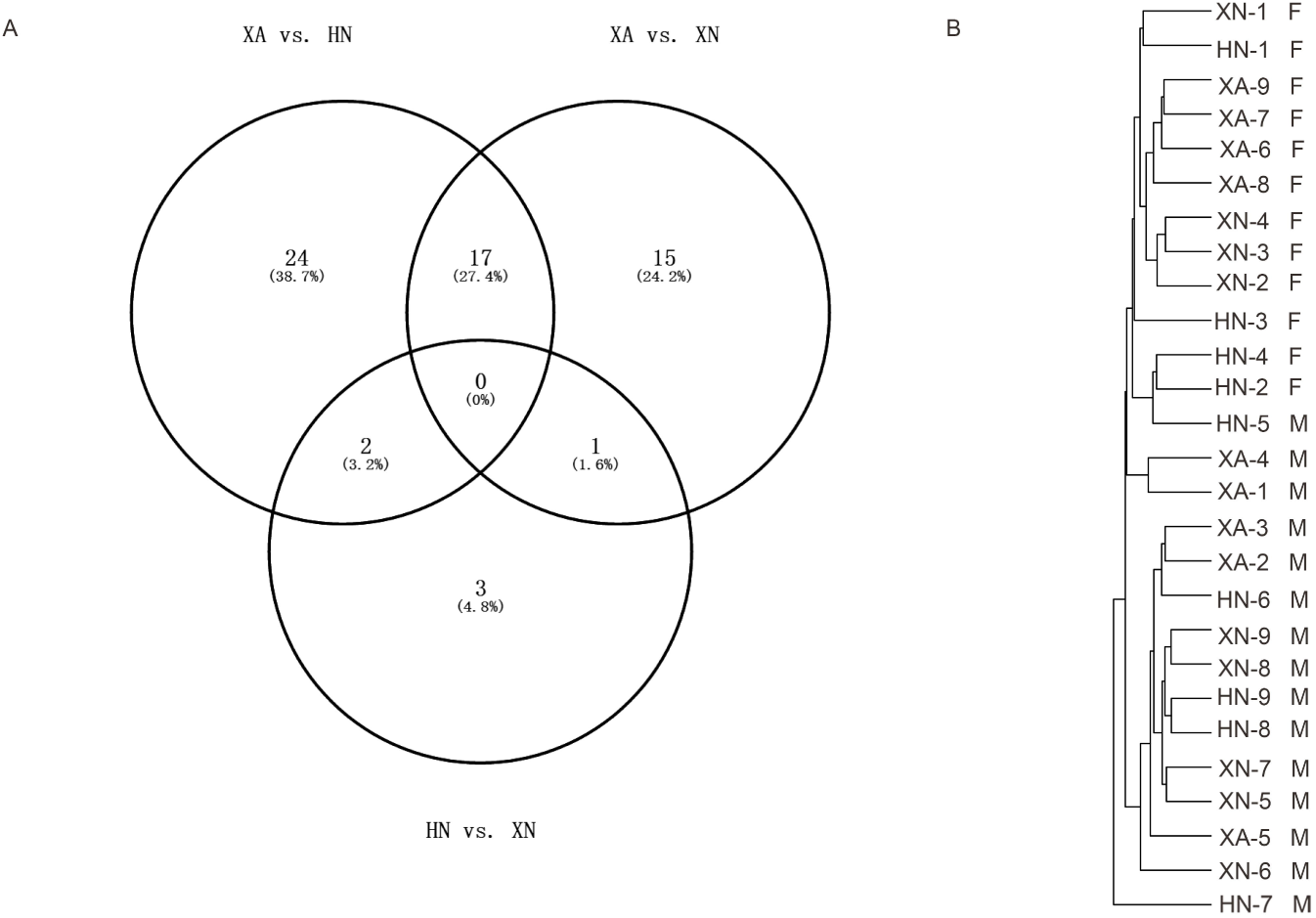
The effects of inter-regional and inter-sex differences on the urine proteome. (A) Veen diagram of the differentially expressed urine proteins (fold change > 2 and p < 0.05) among the different regions. (B) Clustering analysis of all the identified urine proteins from the 27 Urimem samples. HN (Haikou), XA (Xi’an), XN (Xining), M (Male), and F (Female).

Sex is a confounding factor that may influence the urinary proteome. As shown in **Figure 3B**, a hierarchical clustering analysis of all identified proteins was performed to comprehensively define inter-regional and inter-sex differences in the urinary proteome. The results showed that all female samples were clustered into one group and that several samples in the same area could be clustered together. These results suggested that male and female urinary proteomes exhibited different patterns, and inter-regional differences were smaller than inter-sex differences. Between the male and female samples, a total of 53 urine proteins showed more than a 2-fold change and had a p-value < 0.05 (**Table S4**). Interestingly, several sex-specific proteins, such as highly expressed prostate-specific antigen and beta-microseminoprotein, were identified in the male samples.

### Ten disease biomarkers are differentially expressed in the three regions

Of the 62 differentially expressed urine proteins among the three areas, 10 have been reported as disease biomarkers or biomarker candidates in clinical studies (**Table 1**). Peroxiredoxin-3 is involved in cell redox regulation and protects against oxidative stress. Recently, peroxiredoxin-3 was identified as a novel molecular biomarker for hepatocellular carcinoma (Ismail et al., 2015; Shi et al., 2014). Catalase protects cells from the toxic effects of hydrogen peroxide and is involved in oxidative damage. Catalase was reported as a candidate biomarker for colorectal cancer (Maffei et al., 2011), and this oxidative stress biomarker was also upregulated in the urine samples of diabetic patients (Kurutas et al., 2015). Fibrinogen beta chain has a major function in hemostasis, with increased expression reported in the urine samples of patients with deep vein thrombosis and pulmonary embolism (Klovaite et al., 2013; von Zur Muhlen et al., 2016). Alpha-1-acid glycoprotein 2 (Orosomucoid-2) is a candidate plasma biomarker for cholangiocarcinoma (Rucksaken et al., 2017), and is differentially expressed in the urine of pre-eclampsia and diabetes patients (Kronborg et al., 2007; Narita et al., 2004). Dihydropyrimidinase, neuropilin-1 and lysosome membrane protein 2 are disease biomarkers for cancers (Tse et al., 2017; Vasiljevic et al., 2014). Epidermal growth factor receptor kinase substrate 8-like protein 2 and syntenin-1 regulate migration, growth, proliferation, and cell cycle progression in a variety of cancers. These two proteins were reported to be differentially expressed in the urine samples of bladder cancer patients and to be potential cancer biomarkers (Smalley et al., 2008). N(4)-(beta-N-acetylglucosaminyl)-L-asparaginase (AGA) was reported to be increased in the urine sample of prostatitis patients compared with healthy volunteers (Vermassen et al., 2015). Thus, in multi-center biomarker studies, it will be difficult to definitively determine whether potential disease biomarkers are truly associated with disease states or regional differences.

**Table 1.** The effects of regional differences on disease biomarkers

| **ID** | **Protein name** | **MW** | **Ratio** | **P-value** | **Regional difference** | **Biomarker use** |
| --- | --- | --- | --- | --- | --- | --- |
| P30048 | Peroxiredoxin-3 | 27 kDa | 7.70 | 0.038 | Haikou / Xining | Liver cancer |
| P04040 | Catalase | 60 kDa | 4.14 | 0.015 | Haikou / Xining | Colorectal cancer |
| P02675 | Fibrinogen beta chain | 56 kDa | 3.69 | < 0.001 | Xi'an / Xining | Bladder cancer and deep vein thrombosis (urine) |
| P19652 | Alpha-1-acid glycoprotein 2 | 24 kDa | 0.37<br>3.02 | 0.019<br>0.009 | Xi'an / Haikou<br>Haikou / Xining | Cholangiocarcinoma;<br>pre-eclampsia and diabetes (urine) |
| Q14117 | Dihydropyrimidinase | 57 kDa | 2.76<br>2.07 | 0.009<br>0.033 | Xi'an / Haikou<br>Xi'an / Xining | Prostate cancer and colon cancer |
| O14786 | Neuropilin-1 | 103kDa | 2.09 | 0.040 | Xi'an / Haikou | Colorectal Cancer |
| Q14108 | Lysosome membrane protein 2 | 54 kDa | 2.08 | 0.009 | Xi'an / Haikou | Oral Cancer |
| Q9H6S3 | Epidermal growth factor receptor kinase substrate 8-like protein 2 | 81 kDa | 2.12 | 0.021 | Xi'an / Haikou | Bladder cancer (urine) |
| O00560 | Syntenin-1 | 32 kDa | 0.47 | 0.002 | Xi'an / Xining | Renal cell carcinoma (urine) |
| P20933 | N(4)-(beta-N-acetylglucosaminy)-L-asparaginase | 37 kDa | 2.11 | 0.017 | Xi'an / Haikou | Prostatitis (urine) |

2 Footnote: MW-molecular weight

### Functional analysis of differentially expressed proteins

To analyze the functions of the differentially expressed urine proteins, DAVID was used. Forty-three differential proteins between Xi’an and Haikou and thirty-three differential proteins between Xi’an and Xining were uploaded into the database respectively. The most frequently represented categories were biological processes, molecular functions and pathways. The results showed that the main biological processes were the oxidation-reduction process, glycolytic process and reactive oxygen species metabolic process. The main molecular functions included oxidoreductase activity and hydrolase activity, and main biological pathways included metabolic pathways and glycolysis/gluconeogenesis.

## Discussion

In our present study, we investigated whether regional differences can significantly influence the human urinary proteome among different regions. Comparative proteomics analysis of Urimem samples from three areas in China was performed using label-free quantification. Our results indicate that a total of 62 urine proteins were differentially expressed among Xi’an, Hainan and Xining. Of which, 10 proteins were found to be disease biomarkers or biomarker candidates identified in previous clinical studies.

As shown in Figure 2, several blood biochemical indicators were significantly different in the blood of the participants from the three regions, which was consistent with the results of several previous studies. For example, the serum TP and ALB of a healthy population in Xi’an were significantly different from those of some other cities in China (Yin, 2008). Another study suggested that several serum indicators changed with increasing elevation in four areas of Yunnan Province, including BUN, Cr, and uric acid (UA) (Zi, 2014). In another recent study, significant differences in serum immunological indexes and the coagulation index of healthy people between Xi’an and Xining were reported, including a higher expression of IgG, IgM and IgA in Xining than in Xi’an (Gu, 2016). Our results in Table S1 are consistent with this study. Because blood and Urimem samples come from the same participants, and the urinary proteome contains a number of plasma proteins, changes in the blood composition are likely to have a potential impact on the urinary proteome.

In this study, a total of 62 differentially expressed urine proteins were identified among the three areas. The results also suggested that samples in Xi’an were significantly different from samples in the other two cities. Because sex has been reported to influence the urine proteome (Guo et al., 2015), we compared the effects of inter-regional and inter-sex differences on the urinary proteome. A total of 53 differentially expressed urine proteins were identified between the male and female samples. As shown in Figure 3B, the result of the unsupervised clustering analysis indicated that the inter-regional differences were smaller than the inter-gender differences. Previous studies demonstrated that some intrinsic factors (such as sex and age) and extrinsic influences (such as diet, exercise, and environmental factors) may influence the urine proteome, and intrinsic factors likely cause more significant changes to the urine proteome than extrinsic influences (Wu and Gao, 2015). Regional differences are attributable to genetic background, food consumption, lifestyle and environmental factors. In this study, the participants were matched by age, sex and ethnicity. Thus, regional differences in our study were likely due to extrinsic influences, and the result of the clustering analysis is reasonable.

In the present study, Urimem was used to preserve the urine proteins. The system of preserving urinary proteins on membranes using a vacuum filter bottle was first described in our previous study (Jia et al., 2014). The feasibility and validity of Urimem were proven in flow up studies (Qin and Gao, 2015; Wang et al., 2014a; Wang et al., 2014b). The findings showed that for protein absorption, NC membrane had a similar reproducibility to that of PVDF membranes. The proteome coverage of urinary proteins absorbed onto NC membranes has also been evaluated using LC-MS/MS analysis (Wang et al., 2014b). Recently, membrane-based urine protein absorption using a syringe-push system was also described as a simple rapid method for preparing urine for proteomics analysis (Chutipongtanate et al., 2015). These studies demonstrated that dry membranes can be stored at room temperature for an extended period of time are stable for further proteomic analysis. The long-term stability of membranes with dried adsorbed urine proteins allows specimen collection from areas without freezers and overcomes sample shipping problems, such as protein stability during sample shipping and shipping expenses due to weight. Membrane-based methods for urine protein preservation are quick and suitable for batch sample processing in large-scale clinical studies. Additionally, membranes with different properties can be used to selectively retain different substances from urine, such as DNA, metabolites and microRNAs (Zhang et al., 2015). Massive sample storage (biobanking) in a simple and economical manner is the foundation of future large-scale clinical precision medicine studies, and this membrane-based method may become one good option for urine sample processing and storage (Gao, 2015). Urimem is a membrane that can preserve urinary proteins simply and economically. This is the first application of Urimem in epidemiological studies. Urimem is a timesaving and suitable tool for large-scale and rapid sample preservation, especially in places without centrifugation or refrigeration facilities. The extensive use of Urimem in and out of hospitals will accelerate biobank development and future urine biomarker studies using proteomics approaches.

In this study, regional differences in normal urine proteome were evaluated in the urine samples of urban residents from three Chinese regions. Some urinary proteins were differentially expressed in different regions, and several proteins were reported in previous clinical studies to be disease biomarkers. The potential application of our findings in biomarker discovery is that regional differences should be taken into consideration in biomarker identification and validation, particularly in multi-region and multi-center clinical studies. The object of this study was to assess the presence of significant differences in the urinary proteomes of individuals from different regions. Due to the low throughput of MS, only 27 Urimem samples are used for the proteomics analysis, and the results should be confirmed in future studies with a larger sample size and additional populations in other regions.

## Materials and Methods

### Participants

The China National Health Survey (CNHS) was conducted to evaluate physiological constants and health conditions of the Chinese population. Research on the normal urine proteome of the healthy Chinese population is a subproject of CNHS. The study was approved by the local Ethics Committee of the Institute of Basic Medical Sciences, Chinese Academy of Medical Sciences (No. 029-2013), and signed informed consent was obtained from all participants. A total of 754 healthy volunteers between 10 and 80 years of age were enrolled in this study. Urine samples from 283 healthy urban residents (164 males and 119 females) in Haikou, Hainan Province, China, were collected on December 27-29, 2013; urine samples from 255 healthy urban residents (121 males and 134 females) in Xi an, Shaanxi Province, China, were collected on July 8-10, 2014; and urine samples from 216 healthy urban residents (128 males and 88 females) in Xining, Qinghai Province, China, were collected on August 15-17, 2015.

### Data collection

A questionnaire was administered to obtain information from the participants, including sex, age, ethnicity, personal medical history, education level, physical activity, smoking status and drinking status.

Blood pressure was measured on seated participants using an Omron digital blood pressure measuring device. After a donor had fasted overnight, a venous blood sample was collected at the time of the interview. Blood samples were processed within 4 h of collection, shipped to a laboratory in Beijing, and stored at −80°C until analysis.

Blood biochemical and immune indicators, including total cholesterol (TC), triglycerides (TG), high-density lipoprotein cholesterol (HDL-C), low-density lipoprotein cholesterol (LDL-C), immunoglobulin G (IgG), immunoglobulin A (IgA), immunoglobulin M (IgM), alanine aminotransferase (ALT), serum total protein (TP), serum albumin (Alb), the ratio of serum albumin to globulin (AG), gamma glutamyl transferase (GGT), alkaline phosphatase (ALP), aspartate aminotransferase (AST), calcium (Ca), phosphorous (P), creatinine, urea nitrogen, fasting venous blood glucose (Glu) and uric acid (UA), were assessed at the Peking Union Medical College Hospital and Chinese PLA General Hospital.

### Urinary protein preservation on a membrane

Approximately 50-100 mL of morning midstream urine was collected from each healthy participant. The urinary protein concentration was determined using routine urine tests. The urinary proteins were preserved on nitrocellulose (NC) membranes using a filtering device as previously reported (Jia et al., 2014). Briefly, after urine collection, 20 mL of each urine sample was mixed with 10 mL of phosphate buffer (1 M Na_2_HPO_4_, 1 M NaH_2_PO_4_, pH = 6). The diluted urine sample was then added to a vacuum filter bottle (10-cm^2^ filter area). In this device, a polyvinylidene fluoride (PVDF) membrane (0.45 μm HV, Millipore, Tullagreen, Carrigtwohill, Ireland) with ultra-low protein-binding capacity was used to filter cell debris from the urine. Another 0.2-μm NC membrane (GE Healthcare, Freiburg im Breisgau, Germany) was used to adsorb urine proteins. Three sheets of filter paper were wetted with pure water and placed below the NC membrane. The vacuum filter bottle was connected to a vacuum pump, and the urine in the bottle passed through the PVDF membrane, while the urine protein in the flow-through was saved on the NC membrane. Then, the protein-bound NC membrane was dried with a hair dryer at room temperature and sealed in a sterile plastic bag with a label. The medical record number, sex, age, date of urine collection and blood pressure were recorded on the label. Urine samples from each participant were preserved on 2 or 3 NC membranes (Urimem).

### Elution of urinary proteins from the membranes

To avoid potential interference from confounding factors, Urimem membranes from healthy urban Han Chinese adults aged between 30 and 36 with no chronic or metabolic diseases (hypertension, diabetes, hepatitis and cancers) and no drug treatment within 2 weeks were selected. Additionally, blood biochemical indicators from each participant were reviewed to select healthy individuals whose blood indicators fell within the reference ranges for Chinese populations. To rule out potential kidney disease, participants without severe proteinuria were selected after a routine urine protein test. Finally, 9 participants (5 males and 4 females) from each geographic area were randomly selected for further proteome analysis.

Urinary proteins were eluted from the membranes according to a previously described method (Qin and Gao, 2015). The NC membrane with bound proteins was cut into small pieces and placed in a 2-mL Eppendorf tube. Then, 1.7 mL of acetone and 0.2 mL of 5% NH_4_HCO_3_ were added to the tube and vigorously vortexed for 30 s. The tubes were heated at 55°C for 60 min and were strongly vortexed for 30 s every 20 min. After the membrane was fully dissolved, the liquid was gently mixed using a rotary mixer at 4°C for 2 h to precipitate the protein. Then, the mixture was centrifuged at 12,000 x g for 15 min at 18°C. After the supernatant was removed, the tube was placed upside down on filter paper for 5 min at room temperature. The precipitate was re-suspended in lysis buffer (8 M urea, 2 M thiourea, 50 mM Tris and 25 mM DTT), and the protein concentration was determined using the Bradford method.

### Tryptic digestion and LC-MS/MS analysis

A mass of 200 μg of urinary proteins eluted from each Urimem sample was digested with trypsin (Trypsin Gold, mass spectrometry grade, Promega) using the filter-aided sample preparation method with 10-kDa filtration devices (Pall, NY, USA) at 37°C overnight. The peptide mixtures were then desalted using Oasis HLB cartridges (Waters, Milford, MA, USA), dried by vacuum evaporation and re-dissolved in 0.1% formic acid.

For the LC-MS/MS analysis, 1 μg of peptides from each individual sample was loaded onto a trap column (75 μm × 2 cm, 3 mm, C18, 100 Å) and was separated on a reverse-phase analytical column (50 μm × 150 mm, 2 μm, C18, 100 Å) using the EASY-nLC 1200 HPLC system (Thermo Fisher Scientific). The elution for the analytical column was performed over 60 min at a flow rate of 300 μL/min. Then, the eluted peptides were analyzed using an Orbitrap Fusion Lumos Tribrid mass spectrometer (Thermo Fisher Scientific, Waltham, MA, USA). MS data were acquired in the high-sensitivity mode using the following parameters: data-dependent MS/MS scans per full scan with top-speed mode (3 s), MS scans at a resolution of 120,000 and MS/MS scans at a resolution of 30,000 in Orbitrap, 30% HCD collision energy, charge-state screening (+2 to +7), dynamic exclusion (exclusion duration 30 s) and a maximum injection time of 45 ms.

### Label-free quantification

Label-free quantitation of the proteomic data was performed using Progenesis LC-MS software (version 4.1, Nonlinear, Newcastle upon Tyne, UK) according to the procedure described by Hauck et al. (Hauck et al., 2010). Twenty-seven sample features were automatically aligned according to their retention times to maximally overlay all the two-dimensional feature maps (m/z and retention time). Features with charge states of +2 to +4 were selected for further analysis. After alignment and feature exclusion, the samples were divided into three groups (Haikou, Xi’an and Xining) using a between-subject design, and the raw abundances of all features were normalized. Normalization corrects for factors resulting from experimental variation and was automatically calculated over all features in all samples.

All MS/MS spectra were exported from the progenesis software, and the data were searched against the SwissProt Human database (08/2013, containing 20,267 sequences) using Mascot software (version 2.5.1, Matrix Science, London, UK). The parent ion tolerance was set at 10 ppm, and the fragment ion mass tolerance was set to 0.05 Da. A maximum of two missed cleavage sites in the trypsin digestion was allowed. Carbamidomethylation of cysteines was set as a fixed modification, and the oxidation of methionine was considered a variable modification. For quantification, all peptides with an ion score ≥ 30 and p < 0.01 were included, and the total cumulative abundance of a specific protein was calculated by summing the abundances of all peptides allocated to it. Calculations of the protein p value (one-way ANOVA) were then performed on the sum of the normalized abundances across all runs. ANOVA values of p < 0.05 and additionally regulation of ≥ 2-fold were regarded as significant for all further results.

### Functional and pathway analyses

Functional annotation of the differentially expressed proteins among the three areas was analyzed using DAVID (Huang da et al., 2009). Protein classification was performed using gene ontology (GO) analysis for molecular functions and biological processes. Differentially expressed proteins were also uploaded into the Kyoto Encyclopedia of Genes and Genomes (KEGG) for pathway analysis.

### Statistical analysis

Statistical analyses were performed with GraphPad Prism version 7.0 (GraphPad, San Diego, CA, USA). The clinical characteristics data are expressed as the means ± SDs. Comparisons among the three groups were conducted using a one-way ANOVA followed by multiple comparisons with the least significant differences (LSD) test. Group differences resulting in p-values < 0.05 were considered statistically significant.

### Compliance and ethics

The study was approved by the local Ethics Committee of the Institute of Basic Medical Sciences, Chinese Academy of Medical Sciences (No. 029-2013), and signed informed consent was obtained from all participants. The authors declare that they have no competing interests.

## Acknowledgements

This work was supported by the National Key Research and Development Program of China (No. 2016 YFC 1306300), the National Basic Research Program of China (973 Program) (No. 2013CB530805, No. 2013FY114100), the Beijing Natural Science Foundation (No. 7173264, 7172076), the Fundamental Research Funds for the Central Universities (No.10300-31042110), and Beijing Normal University (No. 11100704). We thank Wei Sun (Core Facility of Instrument, Institute of Basic Medical Sciences, Chinese Academy of Medical Sciences) for the technical assistance with the proteomic data analysis.

## Additional files

**Table S1.** Characteristics of 27 participants according to location.

**Table S2**. Total urine proteins identified from the Urimem samples using label-free identification

**Table S3**. Differentially expressed urine proteins among the three areas.

**Table S4**. Gender differences in the urine proteome.

**Figure S1.** Hierarchical clustering analysis of the differentially expressed proteins among the three areas. Heatmap of differentially expressed urine proteins analyzed via ANOVA (p < 0.05). HN (Haikou City), XA (Xi’an City), and XN (Xining City).

**The supporting information (Table S2-S4 and Figure S1) are uploaded at https://figshare.com/s/130588f37a1a79846976**.

